# Illuminating women’s hidden contribution to the foundation of theoretical population genetics

**DOI:** 10.1101/360933

**Authors:** Samantha Kristin Dung, Andrea López, Ezequiel Lopez Barragan, Rochelle-Jan Reyes, Ricky Thu, Edgar Castellanos, Francisca Catalan, Emilia Huerta Sanchez, Rori V. Rohlfs

**Affiliations:** San Francisco State University, Department of Biology, San Francisco, CA 94132; University of California, Merced, Division of Molecular and Cell Biology, Merced, CA 95340

**Keywords:** acknowledged programmer, authorship, computational biology, population genetics, women in science

## Abstract

Plentiful evidence shows an historic and continuing gender gap in participation and success in scientific research. However, less attention has been directed at clarifying obscured contributions of women to science. The lack of visible women role models (particularly in computational fields) contributes to a reduced sense of belonging and retention among women. We seek to counteract this cycle by illuminating the contribution of women programmers to the foundation of our own fields—population and evolutionary genetics. We consider past ‘acknowledged programmers’ (APs), who developed, ran, and sometimes analyzed the results of early computer programs. Due to authorship norms at the time, these programmers were credited in the acknowledgments sections of manuscripts, rather than being recognized as authors. For example, one acknowledgement reads “I thanks Mrs. M. Wu for help with the numerical work, and in particular for computing table I.”. We identified APs in *Theoretical Population Biology* articles published between 1970 and 1990. While only 7% of authors were women, 43% of APs were women. This significant difference (p = 4.0×10^−10^) demonstrates a substantial proportion of women’s contribution to foundational computational population genetics has been unrecognized. The proportion of women APs, as well as number of APs decreased over time. These observations correspond to the masculinization of computer programming, and the shifting of programming responsibilities to individuals credited as authors (likely graduate students). Finally, we note recurrent APs who contributed to several highly-cited manuscripts. We conclude that, while previously overlooked, historically, women have made substantial contributions to computational biology.

---

In the 1970s, leaders in population genetics developed innovative theories and methods to test evolutionary hypotheses [Ewens 1972, Felsenstein 1972, Watterson 1975]. Many of these seminal methods continue to enjoy wide application with current data, some 40 years later. Innovative population geneticists developed these methods despite a lack of data. Especially in the absence of large datasets, computer simulations and numerical approaches were essential to validate these methods. Based on authorship, it seems that this foundational research was conducted by a relatively small number of independent individual scientists, nearly all of whom were men.

In some of the seminal papers from this time, we noticed that non-author computer programmers are thanked in the acknowledgments [Watterson 1975]. While these contributions may have resulted in authorship today, this practice was well within authorship norms of the time. Today, when the historical scientific contributions of women and people of color are being increasingly revealed to popular audiences (*e.g., Hidden Figures*) [Shetterly 2016, Evans 2018], we were curious about the scientific contribution of these “acknowledged programmers” (APs) to population genetics.

## Quantifying the contribution of Acknowledged Programmers

We selected *Theoretical Population Biology* (*TPB*) as our target journal because of its high density of foundational population genetics articles that involved programming. We manually collected the author names, institutional affiliations, acknowledgements text, and APs for all articles published in *TPB* 1970 to 1990. We classified both authors and APs into binary gender categories (men and women, see Supplemental Information).

Cumulatively, over 883 articles, of individuals with classifiable binary gender, significantly more APs were women (43.2%) as compared to authors (7.4%) (Table 1) (two-tailed Fisher exact test, *p* = 4.0×10^−10^). This difference is even more striking when considering the 1970s on their own, when 7.0% of authors were women and 58.6% of APs were women (Supplemental Table 2). In this era, if we consider women APs, the contributions of women are nearly 150% higher than if we only consider authorship.

**Table 1:** Author and acknowledged programmer gender.

|  | Authors | APs |
| --- | --- | --- |
| Female | 80 | 19 |
| Male | 998 | 25 |
| Ambiguous | 164 | 2 |

The acknowledgement of women programmers peaks in the mid 1970s, after which the ratio of women APs decreases significantly (Figure 1, Supplemental Figures 1 and 2, Supplemental Table 3, one-tailed Fisher exact test, *p* = 4.3×10^−3^). This parallels the broader cultural shift which moved computer programming from pink collar work to a respected male-dominated field [Vogel, 2017]. Between the 1970s and 1980s, the practice of acknowledging programmers lost popularity as programming duties were likely transferred to graduate students who received authorship (Supplemental Table 4, two-tailed Fisher exact test, *p* = 0.034) [W. Hill, personal communication, May 2018].

**Figure 1:**
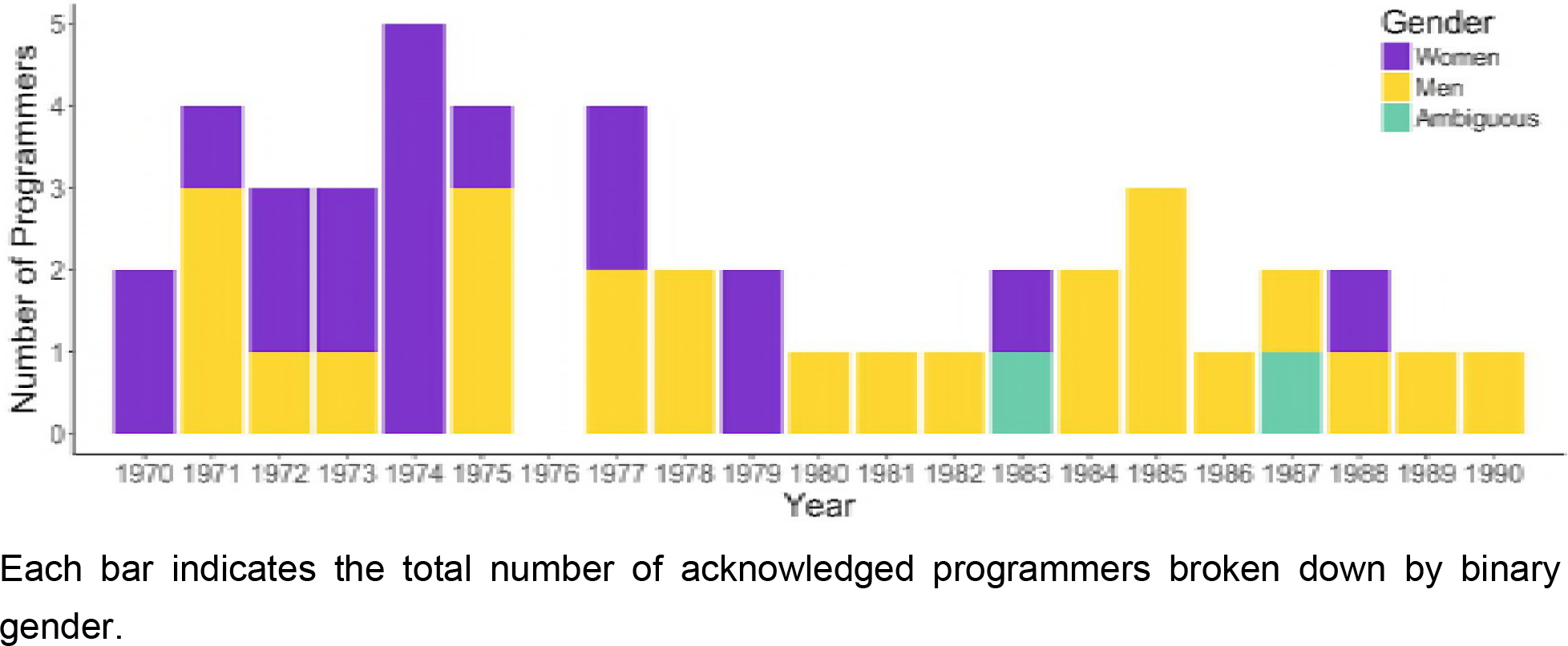
Total number of programmers acknowledged per year in *Theoretical Population Biology*.

## Acknowledged Programmer narratives and contributions

To begin to assess if papers with AP contributors had a disproportionate impact, we compared the number of citations for AP, versus non-AP papers. More high-citation papers (>500 citations) were produced with APs (one-tailed Fisher exact test *p* = 0.055, Supplemental Figure 3, Supplemental Table 5).

In our data, three APs were acknowledged more than once over the years. When Barbara McCann worked as a research assistant at Brown University [Milbank, 1970], she was an AP for two articles in *TPB* (Supplemental Table 1), as well as authoring two papers (Supplemental Table 8). Jennifer Smith was acknowledged for programming and numerical analysis in three articles in *TPB*, as well as at least three additional articles in *Biometrics* when she was a computing assistant at University of Edinburgh (Supplemental Table 6). Lastly, while Margaret Wu was a research assistant in the Department of Mathematics at Monash University, she was acknowledged in two papers in *TPB*, one of which has been cited over 3300 times as it established a widely used estimator of genetic diversity (Supplemental Table 9). She was an AP in at least three additional manuscripts (see Supplemental Methods and Supplemental Table 7). Dr. Wu went on to earn a PhD and hold a faculty position at the University of Melbourne where she developed statistical methods to analyze educational data [Wu, 2011].

The specific technical contribution of an AP likely varied over projects. However, the fact that authors repeatedly chose to work with some APs suggests that these recurrent APs contributed particular expertise. Specifically, in addition to programming and numerical work, Jennifer Smith developed algorithms to carry out verbally specified analyses [W. Hill, personal communication, May 2018]. Margaret Wu performed a variety of statistical work including developing estimators for parameter values, devising algorithms for statistical tasks, and sometimes creating numerical methodology [M. Wu, personal communication, May 2018].

## Scientific contributions and authorship norms

Our retrospective analysis has shed light on the contributions of women to computational genetics research, a field that was drastically skewed towards male authorship. These women’s contributions were previously obscured by being relegated to footnote acknowledgements due to authorship norms. We showed that women’s contribution was substantial when measured by volume (the high proportion of contributions from women APs), as well as by quality when we consider that some women APs were involved in seminal papers and development of cutting edge approaches.

This raises questions about how our current norms of scientific credit may favor certain individuals or groups. For instance, the bibliometric *h* index (*h* such that a scholar authored *h* papers that have been cited at least *h* times) has gained popularity, in part due to its correlation with other indicators of academic success such as National Academy membership or Nobel prize laureateship [Hirsch, 2005]. However, this concordance may reflect that the *h* index is consistent with biases in scientific recognition processes [Kelly and Jennions, 2007]. Current authorship practices also bear examination. In this age of highly collaborative science, authorship can be complex, including non-contributor authors (estimated in 35% of publications in biology), as well as non-author contributors, particularly technicians (estimated in 56% of publications in biology) [Jabbehdari and Walsh, 2017]. Because scientific role (*e.g*., technician, student, PI) is related to social factors (*e.g*., gender, race, class background, nationality), contributions from particular groups will likely remain obscured.

Revealing the scientific contribution of women is particularly important in a sub-field of biology that has been unusually male-dominated, both historically and currently [Telis, 2017]. We wonder if this represents a general trend where women’s contributions in many scientific fields have been relegated to footnote acknowledgments, contributing to a false impression of a lack of participation of women in STEM fields. This historical narrative and our methodological approach, if applied in other scientific fields, can help to positively change the perception of womens’ role in scientific fields and be a motivating force for continued female participation. Because perceiving under-representation in the field can impact a woman’s performance [Steele, 1997, Cheryan *et al*., 2015], providing a counter narrative in and of itself may play a role in improving gender equity.

## Author Contributions

Conceptualization, E.H.S., and R.V.R; Methodology, E.H.S., and R.V.R; Data Curation, E.C., F.C., S.K.D., A.L., E.L.B., R.-J.R., and R.T.; Investigation, S.K.D., A.L., E.L.B., R.-J.R., R.V.R., and R.T.; Writing, S.K.D., E.H.S., A.L., E.L.B., R.-J.R., R.V.R., and R.T.; Supervision, E.H.S., and R.V.R.

## Acknowledgements

This work was funded by SF BUILD NIH grant number 1UL1GM118985 and EHS was supported by NSF grant NSF-DEB-1557151. We would like to thank Michael DeGiorgio, Tracey Heath, Emily Jane McTavish, Rasmus Nielsen, and Pleuni Pennings for constructive comments on an earlier version of this manuscript, as well as Michael Turelli, Jim Harner, and Michael Rose for helping identify the genders of ambiguously named APs. We are also grateful to Bill Hill for information that shaped our research.

## Supplemental Methods

### Gender classification

Classification into binary gender categories was performed for all authors and APs (Supplemental Tables 1 and 9). While the use of these binary categories excludes transgender and gender nonconforming individuals, limiting the accuracy and scope of our analysis, collecting more detailed gender information from these data is practically infeasible. The binary gender analysis presented here is still informative of broad gender dynamics.

When a full, commonly-understood, gender-specific name, or when honorifics (ex: Miss. or Mr.) were provided, gender categorization was straight-forward. However, gender classification was less obvious when provided an individual’s first initial and or a gender-neutral name. To classify an author’s gender, we sought supplementary evidence. For example, by referring a publication, we learned that J. A. Sved is John A. Sved (Sved, 1971; *Kidwell et al., 1977*). Another example is G.L.Yang, whose full name was identified via a book published in the same field by Grace Lo Yang, who had the same university affiliation (Yang, 1972; Cam and Yang 2000). The sources of our evidence (*i.e*., website addresses) are documented in Supplementary Table 9.

### Programmer classification

We identified programmers through key phrases in acknowledgement section such as “ably programming and executing all the computations” (Supplemental Table 1). However, in some cases the acknowledgements were ambiguous about specific technical role, for example, “carrying out the computing” (Supplemental Table 1). In these cases, we evaluated if the study required programming for the task indicated. The vast majority of individuals acknowledged made non-computational contributions. These non-programmers were acknowledged for contributions like “typing the manuscript,” or “helpful comments and financial support”.

### Citation analysis

Between 7 and 12 June, 2018, we recorded the number of citations for each article in the dataset according to Google Scholar (Supplemental Figure 3). We compare the proportion of high-citation papers (those with at least 500 citations) between AP-supported and non-AP manuscripts, finding nearly significantly more high-citation papers among the AP manuscripts (one-tailed Fisher exact test *p* = 0.055, Supplemental Table 5).

### Repeat acknowledged programmers

We researched more articles outside our original *TPB* dataset where repeat APs were acknowledged. As an illustrative example, consider Jennifer Smith, who was acknowledged by William Hill in *TPB*. We used Google Scholar to find articles by “William G. Hill” and “W.G.Hill” between the years 1965 and 1980. After ensuring that the William Hill who authored these newly identified manuscripts had the same affiliation as the original William Hill (Institute of Animal Genetics in Edinburgh or Iowa State University), we examined the acknowledgements section in each resulting article. The articles where Jennifer Smith is acknowledged for programming are documented in Supplemental Table 6. Analogous searches were performed for Barbara McCann and Margaret Wu (Supplemental Tables 7 and 8).

### Changing AP gender ratio

To determine if the gender ratio of APs differed between the 1970s and 1980s, we performed a one-tailed Fisher exact test to determine if the proportion of women APs decreased. There were a total of 373 articles in the 1970-1979 and 364 articles in the 1980-1990. There were 17 female programmers and 12 male programmers acknowledged in the 1970s. The 1980s had two female programmers and 13 male programmers. The proportion of female programmers was significantly lower in the 1980s (*p*=0.00425) compared to the 1970s.

**Supplemental Table 1:**
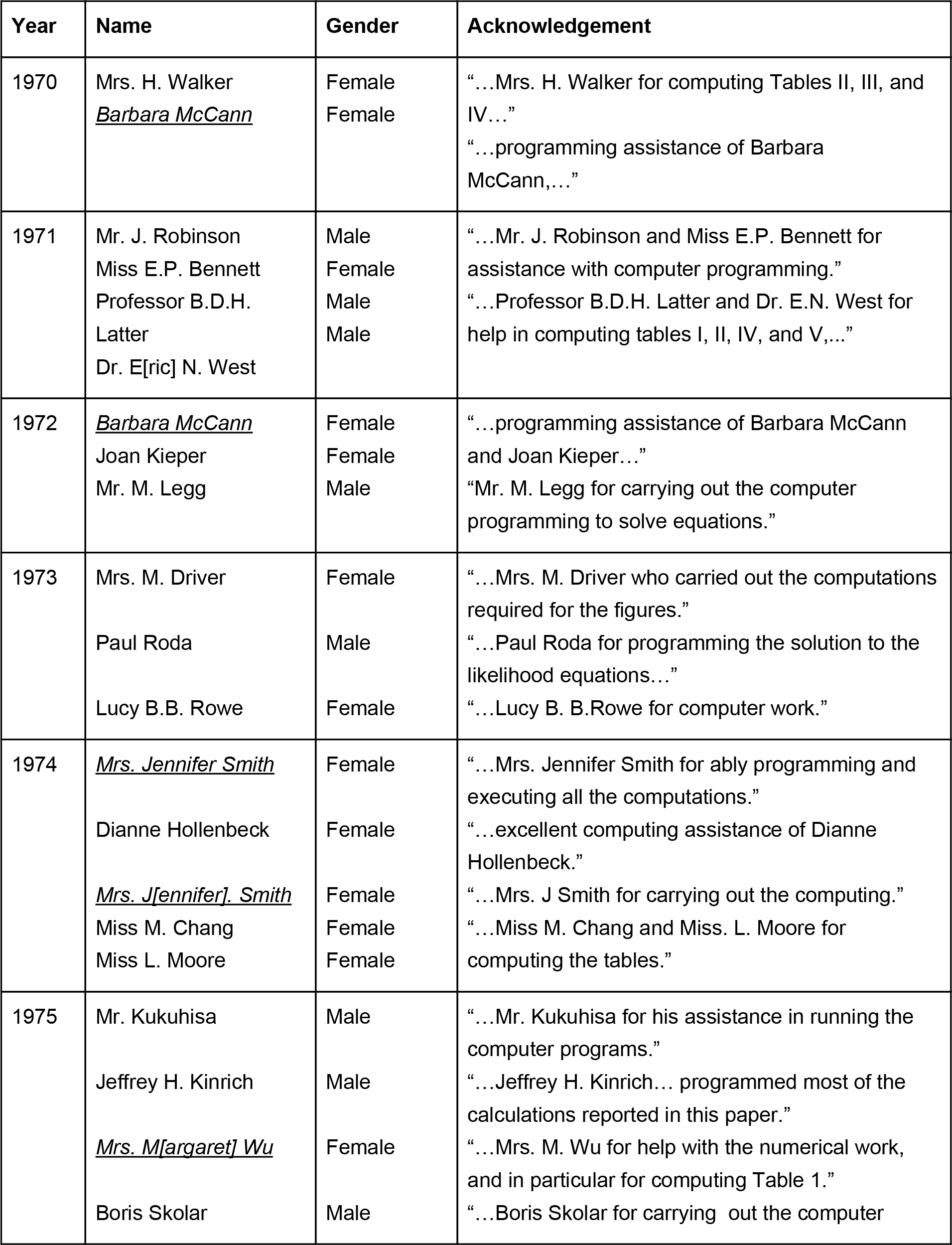
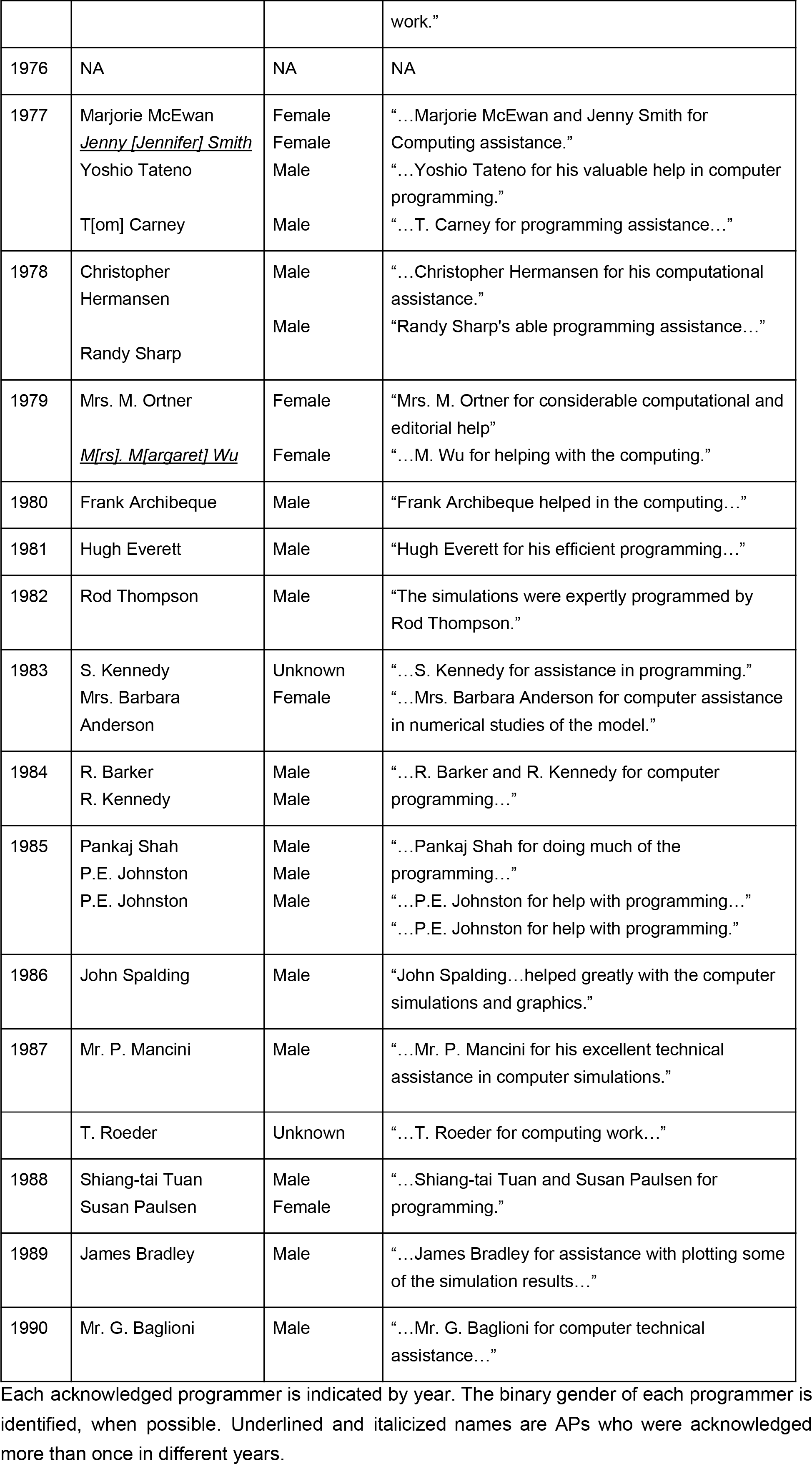
Acknowledged programmers.

**Supplemental Table 2:** Author and acknowledged programmer gender from 1970 through 1979.

|  | Authors | APs |
| --- | --- | --- |
| Female | 38 | 17 |
| Male | 502 | 12 |
| Ambiguous | 24 | 2 |

**Supplemental Table 3:** AP gender ratio over time.

|  |  | Years |  |
| --- | --- | --- | --- |
|  |  | 1970-79 | 1980-1990 |
| Programmer | Female | 17 | 2 |
|  | Male | 12 | 13 |

**Supplemental Table 4:**
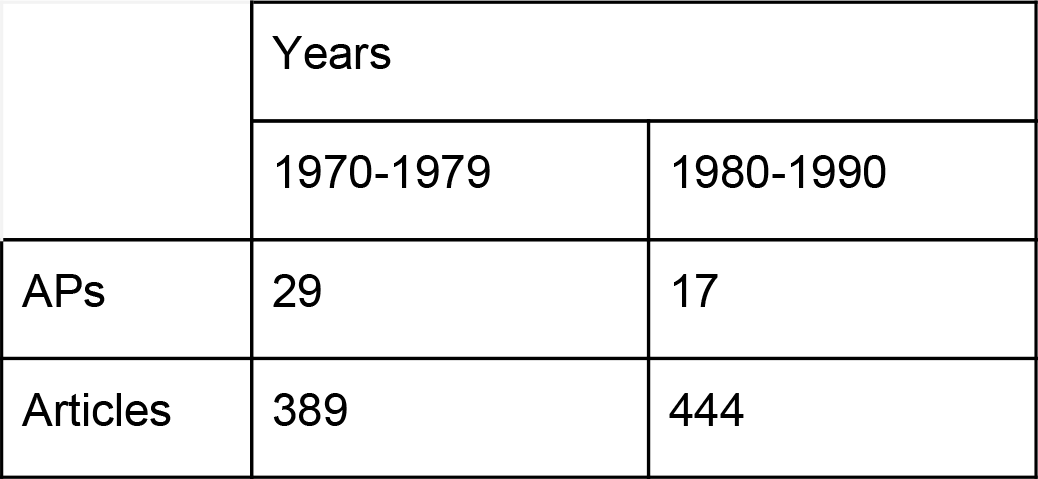
Prevalence of APs over time.

|  | Years |  |
| --- | --- | --- |
|  | 1970-1979 | 1980-1990 |
| APs | 29 | 17 |
| Articles | 389 | 444 |

**Supplemental Table 5:**
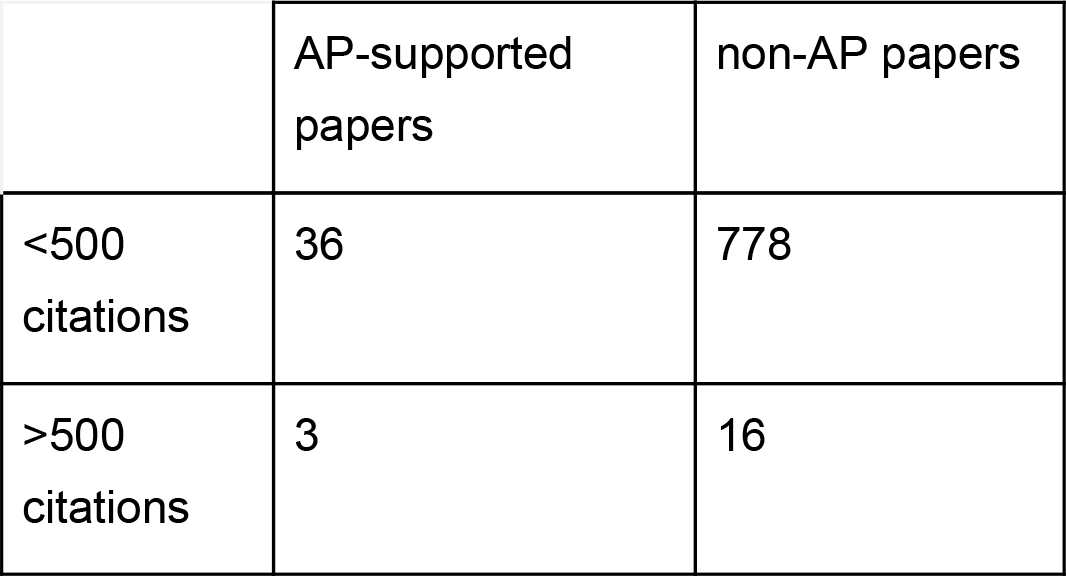
Citations of AP-supported versus non-AP papers.

|  | AP-supported<br>papers | non-AP papers |
| --- | --- | --- |
| <500<br>citations | 36 | 778 |
| >500<br>citations | 3 | 16 |

**Supplemental Table 6:** Acknowledgements for Jennifer Smith.

| Year | Journal | Article Title | Indicative acknowledgments text |
| --- | --- | --- | --- |
| 1975 | *Biometrics* | Tests for Association of Gene Frequencies at Several Loci in Random Mating Diploid Populations | "I am indebted to ... Mrs. Jennifer Smith for programming the analysis on the computer." |
| 1976 | *Biometrics* | Order Statistics of Correlated Variables and Implications in Genetic Selection Programmes | " I am indebted to Jenny Smith for undertaking the numerical analysis..." |
| 1977 | *Biometrics* | Order Statistics of Correlated Variables and Implications in Genetic Selection Programmes. II. Response to Selection | " I am indebted to Jenny Smith for undertaking the numerical analysis." |

This table lists the articles outside of the original *Theoretical Population Biology* dataset where Jennifer Smith was acknowledged for programming. The year, journal, and indicative acknowledgements text are shown.

**Supplemental Table 7:**
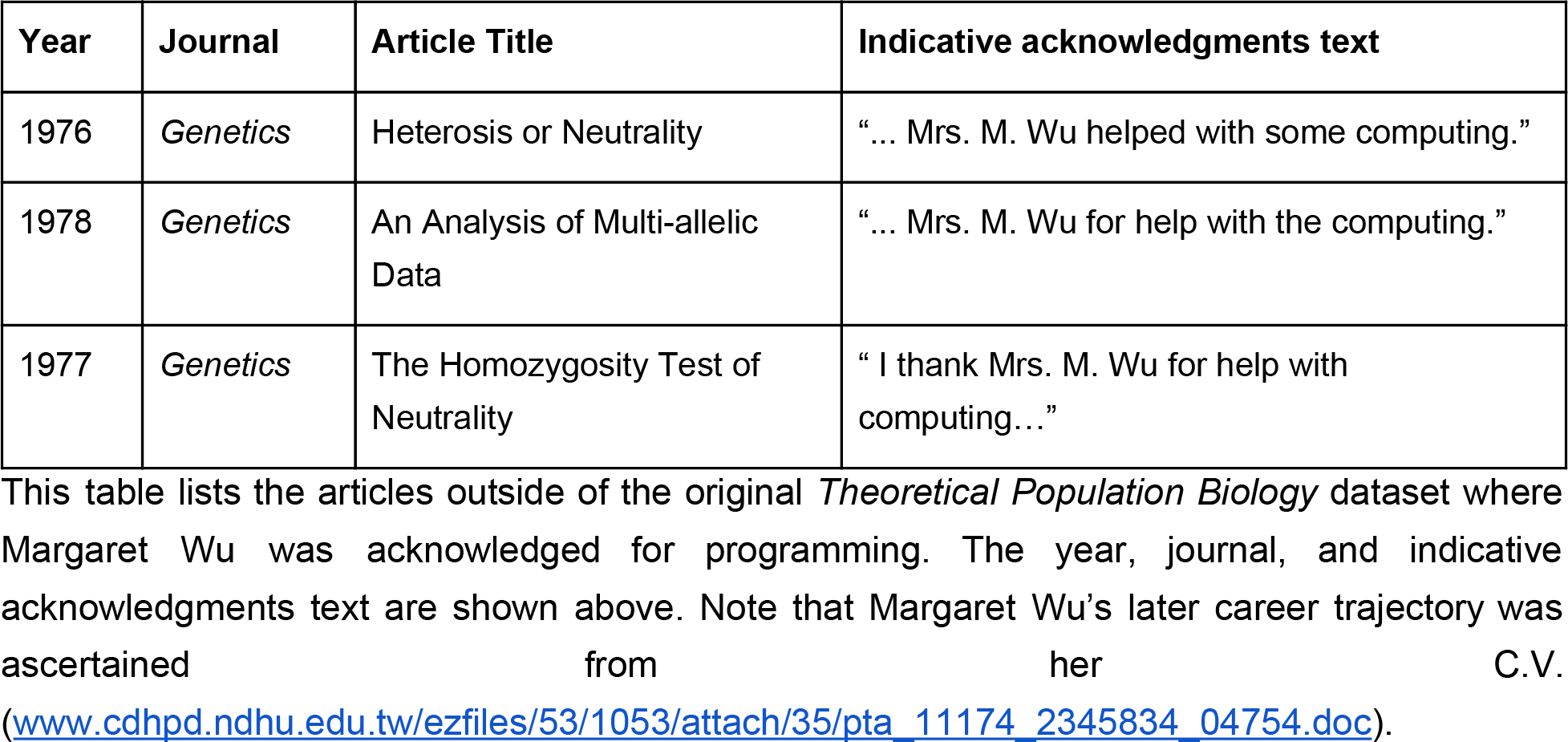
Acknowledgments for Margaret Wu.

**Supplemental Table 8:**
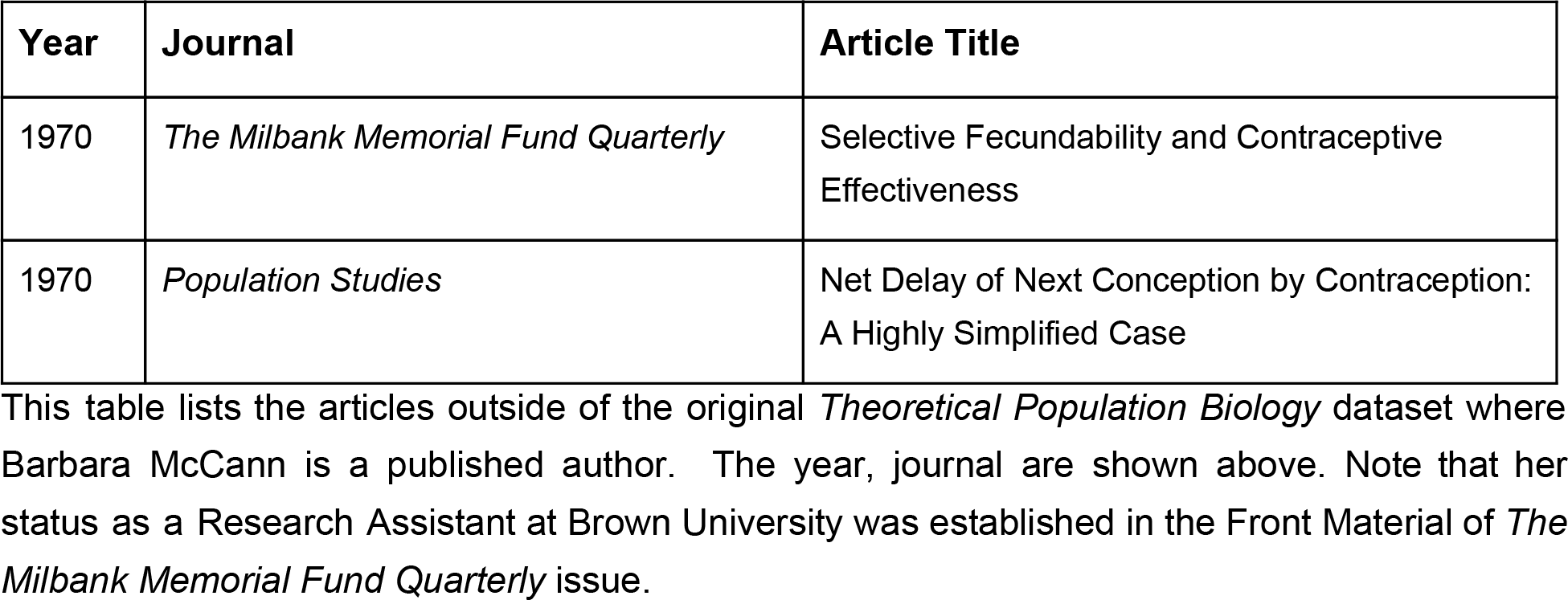
Authorship for Barbara McCann.

**Supplemental Figure 1:**
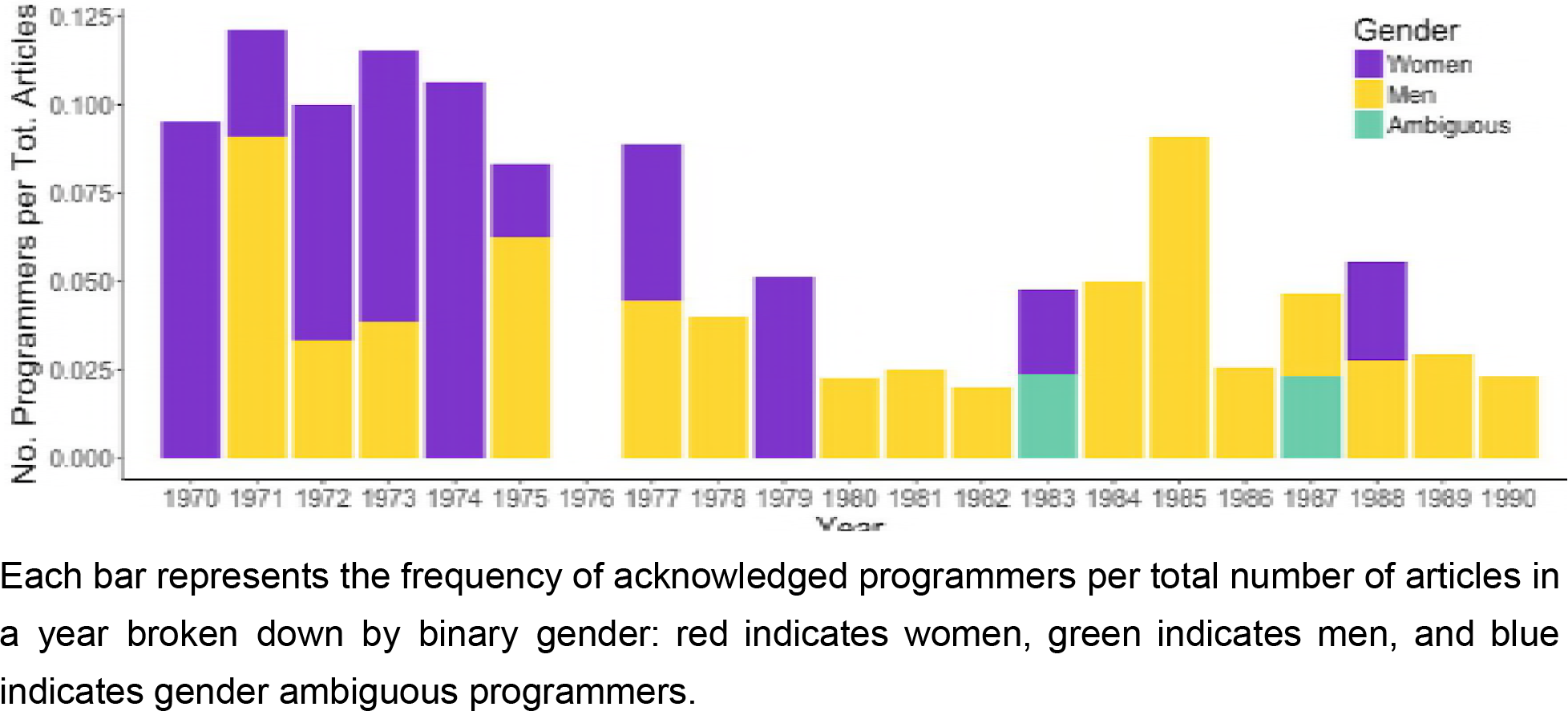
Acknowledged programmers per total number of articles.

**Supplemental Figure 2:**
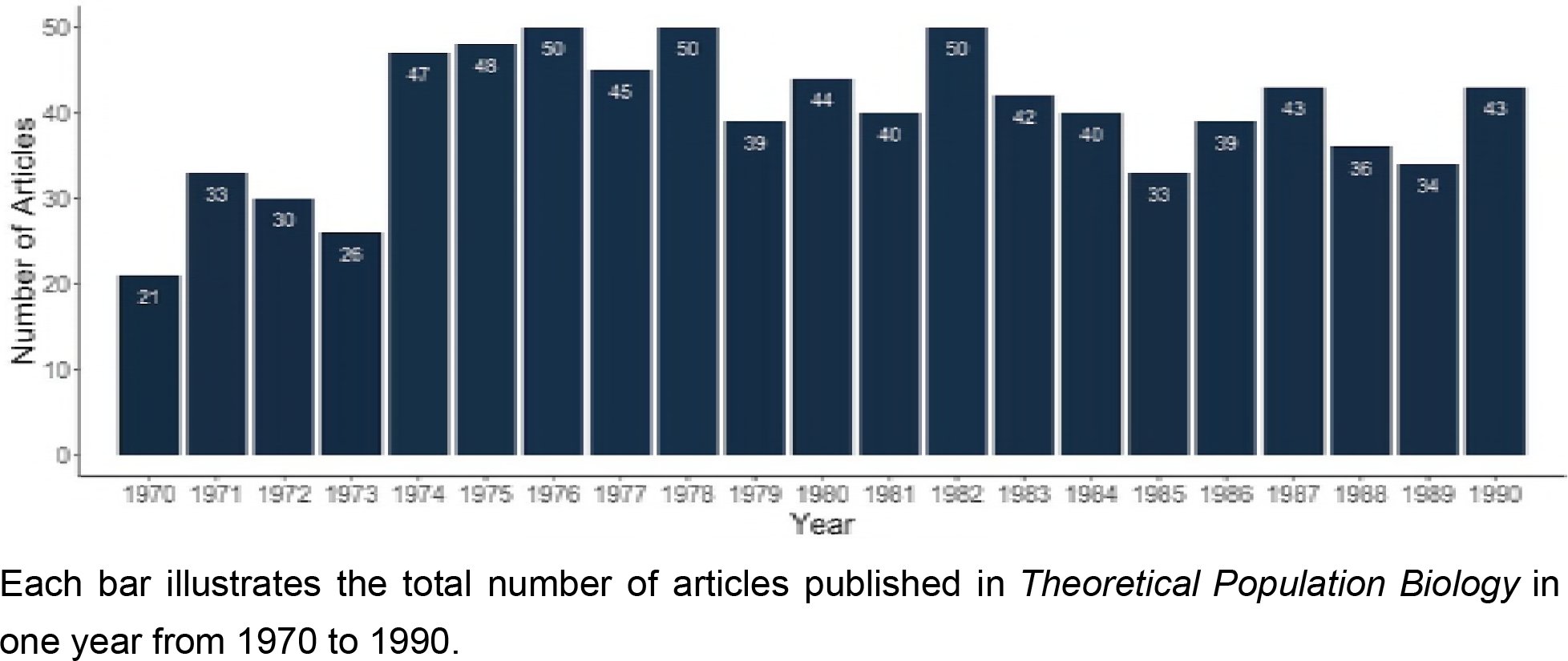
Total number of articles per year in *Theoretical Population Biology*.

**Supplemental Figure 3:**
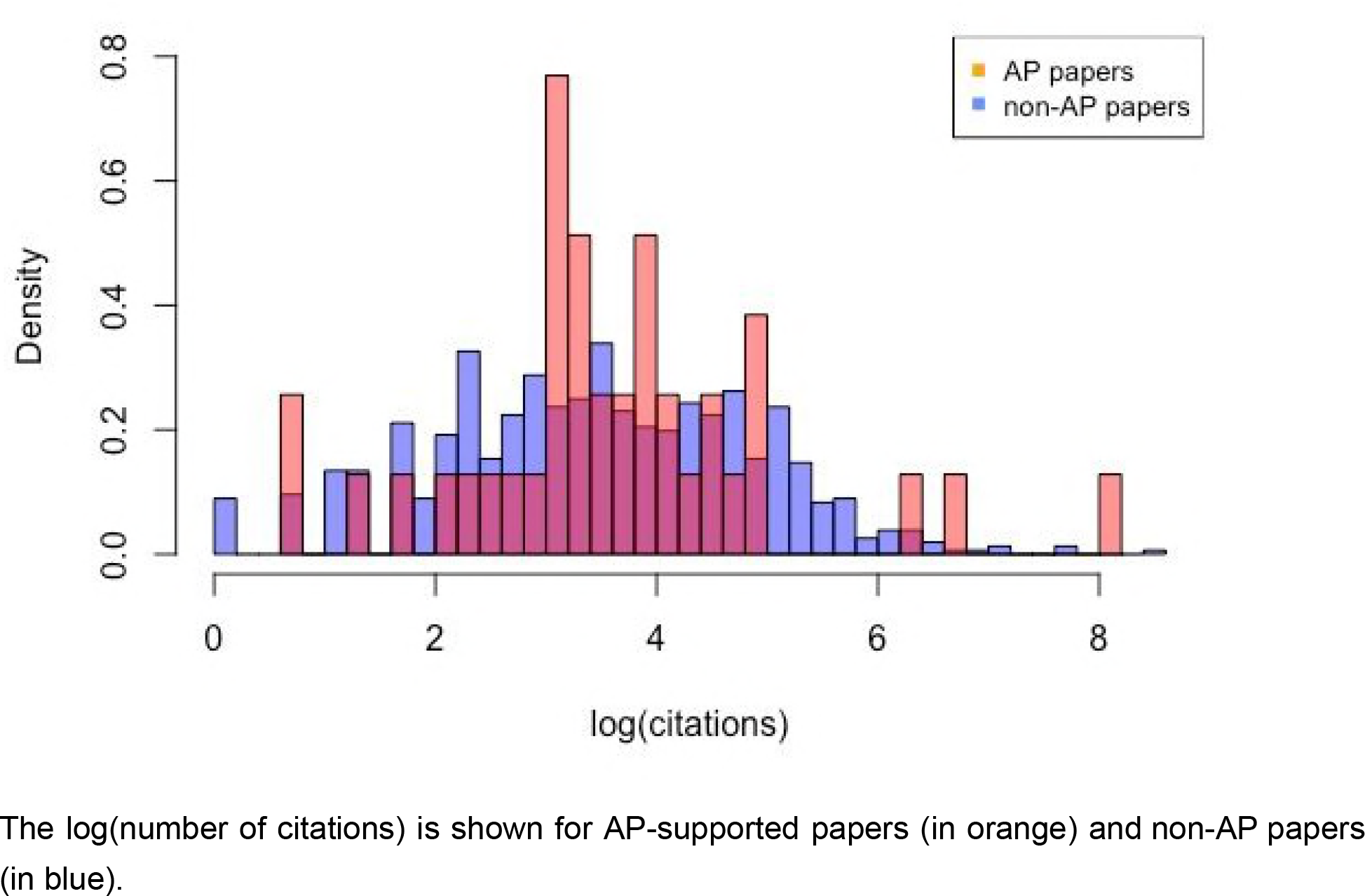
Histogram of number of citations for AP-supported and non-AP papers.

